# Microglia contact cerebral vasculature through gaps between astrocyte endfeet

**DOI:** 10.1101/2024.03.14.585114

**Authors:** Gary P. Morris, Catherine G. Foster, Brad A. Sutherland, Søren Grubb

**Affiliations:** Tasmanian School of Medicine, College of Health and Medicine, University of Tasmania, Hobart, Tasmania, Australia; Center for Developmental Biology and Regenerative Medicine, Seattle Children’s Research Institute, Seattle, WA, USA; Center for Translational Neuromedicine, Faculty of Health Sciences, University of Copenhagen, DK-2200 Copenhagen N, Denmark

**Keywords:** Astrocyte endfeet, capillaries, endothelial cells, microglia, pericytes

## Abstract

The close spatial relationship between microglia and cerebral blood vessels implicates microglia in vascular development, homeostasis and disease. In this study we used the publicly available Cortical MM^3 electron microscopy dataset to systematically investigate microglial interactions with the vasculature. Our analysis revealed that approximately 20% of microglia formed direct contacts with blood vessels through gaps between adjacent astrocyte endfeet. We termed these contact points “plugs”. Plug-forming microglia exhibited closer proximity to blood vessels than non-plug forming microglia and formed multiple plugs, predominantly near the soma, ranging in surface area from ∼0.01 μm^2^ to ∼15 μm^2^. Plugs were enriched at the venule end of the vascular tree and displayed a preference for contacting endothelial cells over pericytes at a ratio of 3:1. In summary, we provide novel insights into the ultrastructural relationship between microglia and the vasculature, laying a foundation for understanding how these contacts contribute to the functional cross-talk between microglia and cells of the vasculature in health and disease.

## Introduction

The vast majority of gray matter astrocytes (∼99.8%) are connected to at least one blood vessel.^1^ These connections, known as astrocyte endfeet, provide almost complete coverage of all cerebral blood vessels.^2^ The area of astrocytic endfeet around a vessel is proportional to vessel diameter, with larger vessels having a greater area of endfeet surrounding them than smaller vessels.^3^ Consistent with this, in larger vessels the astrocyte soma frequently constitute the endfeet.^4^ Furthermore, endfeet surrounding arteries are larger in area than those surrounding veins.^3^ Following injury, astrocyte endfeet coverage is maintained through astrocyte plasticity, but the speed of endfeet recovery declines with age.^5^ The completeness of this coverage also has important functional implications. For example, astrocyte endfeet may control the exchange of water and metabolites between the brain and blood.^2, 3^ They may also play important roles in the regulation of blood flow^6–8^ and glymphatic flow.^9^

Despite astrocyte coverage of the cerebral vasculature being considered almost complete, gaps do exist. One study suggested intercellular and endfoot-endfoot gaps between adjacent astrocytes (also called clefts) are ∼20 nm wide and may make up 0.3% of the area of the brain’s microvessels.^2^ However, the precise width, prevalence and dynamics (i.e if they grow or shrink physiologically^3^) of these gaps remains unclear. It is also not clear if they exist between astrocytes around both large (e.g. arterioles and venules) and small (e.g. capillaries) blood vessels. Their purpose is also unknown, although some have suggested they could provide a sieving function to regulate paravascular solute entry into the brain.^3, 10^

It is also possible these gaps provide entry points for cells of the brain parenchyma to directly contact the cerebral vasculature, which has been observed in a few reports.^2, 4^ Alternatively, cells of the parenchyma may contact the vasculature through discontinuities in individual astrocyte endfeet.^2, 11^ However, it is not yet clear if such discontinuities truly exist, or if they simply reflect astrocyte endfeet becoming so thin that they are difficult to discern, despite still being present.^4^

A growing body of evidence suggests microglia are one such cell that may directly contact the cerebral vasculature without the interference of astrocyte endfeet.^2, 12–20^ Despite the many studies heavily implicating a direct relationship between microglia and the vasculature, the ultrastructural features of microglia-vasculature contact points are poorly defined. In this study we used a publicly available electron microscopy dataset to conduct a formal analysis of the number of microglia directly contacting the vasculature without the structural interference of astrocyte endfeet, and the frequency of these contacts. Furthermore, we assessed where on the vascular tree microglia were more likely to directly contact (e.g. arterioles, venules, or capillaries), which cells microglia were directly interacting with at contact points (e.g. endothelial cells, pericytes etc.) and what the shapes and sizes of these contact points were. Our results lay the foundation for future studies aiming to define the function of microglia-vascular contact.

## Material and Methods

### Defining regions of interest within a public volume electron microscopy dataset

Microglia of the mouse visual cortex were identified within the publicly available Cortical MM^3 dataset (https://www.microns-explorer.org/cortical-mm3).^21^ We aimed to assess microglia-vascular interactions across all levels of the vascular tree including areas containing a penetrating arteriole, capillaries, and an ascending venule. To ensure an unbiased analysis, three adjacent regions of interest (ROI) were created using the box annotation tool in Neuroglancer. Annotations were placed at specific x, y and z coordinates to create ROIs. We created one ROI containing a penetrating arteriole (top left: 189377, 73419, 26850, lower right: 223825, 250250, 23770), one containing an ascending venule (top left: 258374, 73419, 26850, lower right: 292722, 250250, 23770), and one containing the capillary bed between them (top left: 223825, 73419, 26850, lower right: 258374, 250250, 23770, Fig. 1(a-e)).

**Figure 1:**
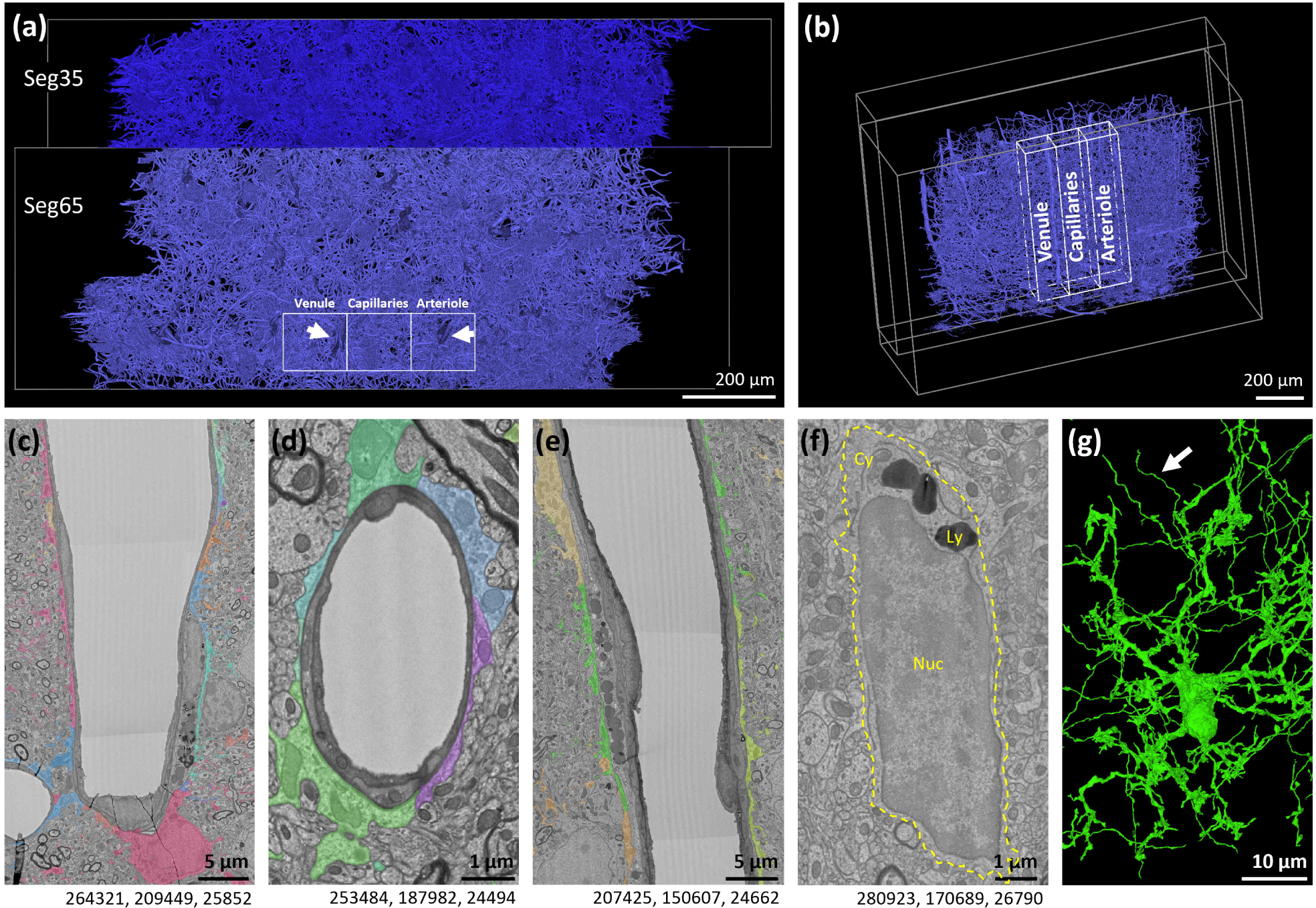
Regions of interest placed around a venule, arteriole and the capillary bed between them. (a) Schematic of our regions of interest within the Cortical MM^3 dataset, looking from the top down. Blood vessel lumens are highlighted in blue. (b) Schematic showing a side view of where regions of interest were placed within the Cortical MM^3 dataset. (c-e) Representative examples of the (c) venule, (d) a capillary and (e) the arteriole within our region of interest. Astrocyte endfeet surrounding each vessel are highlighted in multiple colours. Astrocyte endfeet ultrastructures have been previously described in detail.^4^ (f) Representative example of a microglia nucleus (Nuc), cytoplasm (Cy) and lysosomes (Ly) in the Cortical MM^3 dataset. (g) 3D reconstruction of microglia in (f). White arrow points to an area of the segmentation where a process belonging to a different cell has been picked up alongside the microglia segmentation. Numbers beneath the figures are xyz coordinates from the Cortical MM^3 dataset.

### Identification of different cell types within the Cortical MM^3 dataset

Arteriole and venule starting points were identified using annotations provided by previous studies (https://n9.cl/vid4i ^4, 20^). The colorized 3D segmentation of vasculature in Fig. 6e and Supplementary Fig. 5 were made by annotations in Neuroglancer, extracting coordinates from the JSON file, followed by adding meshes to the resulting point clouds using Meshlab and finally 3D rendering in Blender, as previously described.^27^

Microglia were identified as previously described.^4, 12^ In brief, they were identified by their small soma size compared to other cell bodies, their distinct heterochromatin pattern, thin cytoplasm, prominent dark lysosomes (visible at low magnification), and an elongated irregular shape that often resembles a bean^22, 23^ (Fig. 1(f-g)). Oligodendrocyte progenitor cells (OPCs) are visually similar to microglia, but these were differentiated by the presence of primary cilia in OPCs^24^ (see Supplementary Table 1 for examples), which are absent in microglia.^25^ Endothelial cells were readily identified forming the inner lining of the vasculature^4^ and pericytes were identified on the abluminal surface of endothelial cells on capillaries, with which they share the basement membrane. Both have protruding cell bodies and long processes.^20, 26^ Perivascular macrophages (PVMs) and perivascular fibroblasts (PVFs) were identified as previously described^4^ (see Supplementary Table 2 for a list of PVMs and PVFs in our analysis). Astrocytes were identified by their electron lucent appearance compared to other cell types, and by their extensive processes.^20, 26^ In general, the 3D cell segmentation in the MICrONS dataset assisted with cell identification.

### Annotation of microglia

Specific locations within the Cortical MM^3 dataset were identified using a point annotation tool placed at x, y, z coordinates. These coordinates were used to determine distances between different annotations. Microglia were identified and annotated by navigating each z-plane within the ROIs. The centre point of each microglia was located by finding the middle z-plane of the microglia nucleus, then annotating the centre of the nucleus at that z-plane by eye. The nearest blood vessel to each microglia was determined by the distance between the annotation placed at the centre of the microglia and an annotation placed at the edge of the nearest blood vessel. Blood vessel annotations were placed on the outer edge of the basement membranes.

### Annotation of microglial plugs and fenestrations

Locations where microglia directly contact the vasculature through gaps between two or more adjacent astrocyte endfeet were defined as microglial ‘plugs’. We initially identified plugs with the aid of the coloured segmentation, which enabled us to track microglia processes and the vasculature within greyscale 2D EM images. We note the segmentation is not always accurate (Fig. 1(g))^4, 20^, so the ultrastructure was also carefully scrutinized to confirm if processes belonged to microglia. 12 microglia were unable to be assessed due to poor segmentation. If clear separation could be seen between two close adjacent plugs, they were counted separately. However, if they filled the same gap between astrocytic endfeet, they were counted as one plug. We also assessed where microglia may be contacting the vasculature through gaps in a single astrocyte endfoot, contacts we define as ‘fenestrations’.

Possible plugs and fenestrations were first annotated by GPM, before being reviewed by an independent researcher with prior experience analysing electron microscope datasets (SG). If researchers disagreed or were not convinced about the validity of a plug, it was removed from the dataset. Each plug’s centre point was determined by finding the central point of contact with the vasculature in the middle z-plane and placing an annotation. If the plug process was bifurcated at the middle z-plane, the centre point was placed in the part of the plug closest to the microglia cell soma.

To determine the nearest arteriole or venule to the capillary with plugs, we traced the least number of capillary orders to a larger vessel, always counting from the ascending venule. The few examples of plugs we found 9 orders away from the venule may be closer to a venule (or an arteriole) elsewhere in the brain (Fig. 4(c)), however, because the ROIs selected were near the edge of the EM volume some of the blood vessels extend into territories outside the dataset, so this could not be determined.

### Annotation of microglial centrioles

Microglia centrioles were mostly found in the vicinity of the nucleus by first locating the Golgi apparatus that surrounds them (Fig. 6(b-c)). They are cylindrical and exist in pairs at right angles to each other. In some cases, the centriole pair was located far from the nucleus within one of the larger microglial processes.

### Quantification of plug-adjoining astrocytic endfeet and plug area

The number of astrocyte endfeet surrounding each microglial plug was determined with the aid of the coloured segmentation and by inspecting the ultrastructure. We assessed all z-planes where plugs were present, due to their 3D structures.

To approximate microglial plug area, the number of z-planes each plug spanned was determined. At the middle z-plane, the line annotation tool was used to trace the length of contact. Where it was not possible to assess width at the centre point (due to artefacts e.g. tissue folds), we assessed width at the z-plane directly prior to or after the artefact (whichever plane the contact point was wider on). These two distances were used to calculate the area of an ellipse (area = π x r_1_ x r_2_).

### Quantification of the cells on the vascular that microglial plugs most frequently contact

The identity of vascular cells contacted by each plug was determined. Endothelial and pericyte soma were defined as the region of the cell containing the nucleus, which were readily identifiable in the EM images, and processes were defined as any other parts of the cells.

### In vivo two-photon microscopy of microglia surveying nearby capillary pericytes

All animal procedures were approved by the Animal Ethics Committee, University of Tasmania (A18608 and A23735) and conformed with the Australian National Health and Medical Research Council (NHMRC) Code of Practice for the Care and Use of Animals for Scientific Purposes, 2013 (8th Ed.). All results are reported in accordance with the ARRIVE guidelines.^28^ To investigate microglial surveillance near pericytes, we analysed *in vivo* two-photon microscopy data from a subset of NG2DsRed×CX3CR1^+/GFP^ mice (n = 3) that had been implanted with cranial windows in our previous study^12^. Please refer to our previous publication for extended details including animal husbandry, cranial window implantation and microscopy^12^.

Images taken on day -1, when FITC-dextran had been administered prior to imaging, were used to determine the position of microglia on the capillary tree. A separate set of images of the same microglia and pericytes imaged on day -1, but not previously analysed, were taken on day 14 (see previous publication for imaging timeline^12^). Images were generated using a 20X water immersion lens (NA 1.0, Zeiss Plan-Apo) at 4.0 zoom with 17% laser power and by imaging 1 µm z-step intervals over a total of 25 µm. Each 25 µm stack took ∼45 s to image, after which the loop was immediately restarted until the same region was imaged 10 times. This took a total of ∼7 min 30 s for each region. We imaged a capillary near an arteriole in one male mouse, a capillary in the middle of the capillary bed in one female mouse and a capillary near a venule in one female mouse. Two-photon hyperstacks (3D movies) were corrected for 3D drift using the ImageJ plugin “Correct 3D drift” first in z-direction and then in x-y direction. Subsequently, hyperstacks were volume and surface rendered in Imaris to make the videos.

### Statistical Analysis

Data were processed in Microsoft Excel and statistical analyses were performed using GraphPad Prism 10.1.2 (GraphPad, USA). Data were tested for normality using the D’Agostino & Pearson test. Nuclear-centrosome (NC) lengths of non-plug and plug-forming microglia were not normally distributed and were compared using a Mann–Whitney test. NC lengths in all three ROIs were also not normally distributed so were compared using a Kruskal-Wallis test, followed by a Dunn’s post-hoc test. Overall ANOVA results are reported in the figure legend and results of the post hoc test are represented in the figure and reported in the results text. NC length was compared to nucleus to vessel distance using a Spearman correlation as data was not normally distributed. A p<0.05 was considered statistically significant. Data are presented as frequency distributions, mean ± standard deviation (SD), or median + range (see figure legends for details). The raw data supporting the findings of this study are provided in Supplementary Table 3.

## Results

### The majority of plug-forming microglia are in close proximity to the vasculature

Although evidence has previously been provided that microglia may directly contact the vasculature,^2, 12–19^ there are no prior studies formally quantifying the frequency, number, size or location of microglial contacts with the vasculature using tools with sufficient resolution, such as electron microscopy. We took advantage of the publicly available Cortical MM^3 electron microscopy database^21^ to obtain quantitative information regarding the ultrastructural nature of microglial contacts with the cerebral vasculature (Fig. 1). The dataset provided sufficient resolution to observe microglial contacts with the vasculature that are below the limit of detection of other tools. In addition, the dataset is large enough to determine differences along the entire vascular tree.

We first placed three ROIs encompassing a penetrating arteriole, an ascending venule and the capillary bed between them. Collectively, these ROIs covered ∼3.6% of the total dataset volume. Within these ROIs we found 307 microglia (Fig. 2(a)). 28.7% were in the venule ROI, 39.4% in the capillary ROI and 31.9% in the arteriole ROI. 54.4% of all microglia had their nuclear centre point within 10 µm of a blood vessel, and 17.8% of all microglia were <2.5 µm from a blood vessel. The average distance of microglia to capillaries was 9.7 μm. This data highlights the close spatial relationship between microglia and the cerebral vasculature (Fig. 2(b-d)).

**Figure 2:**
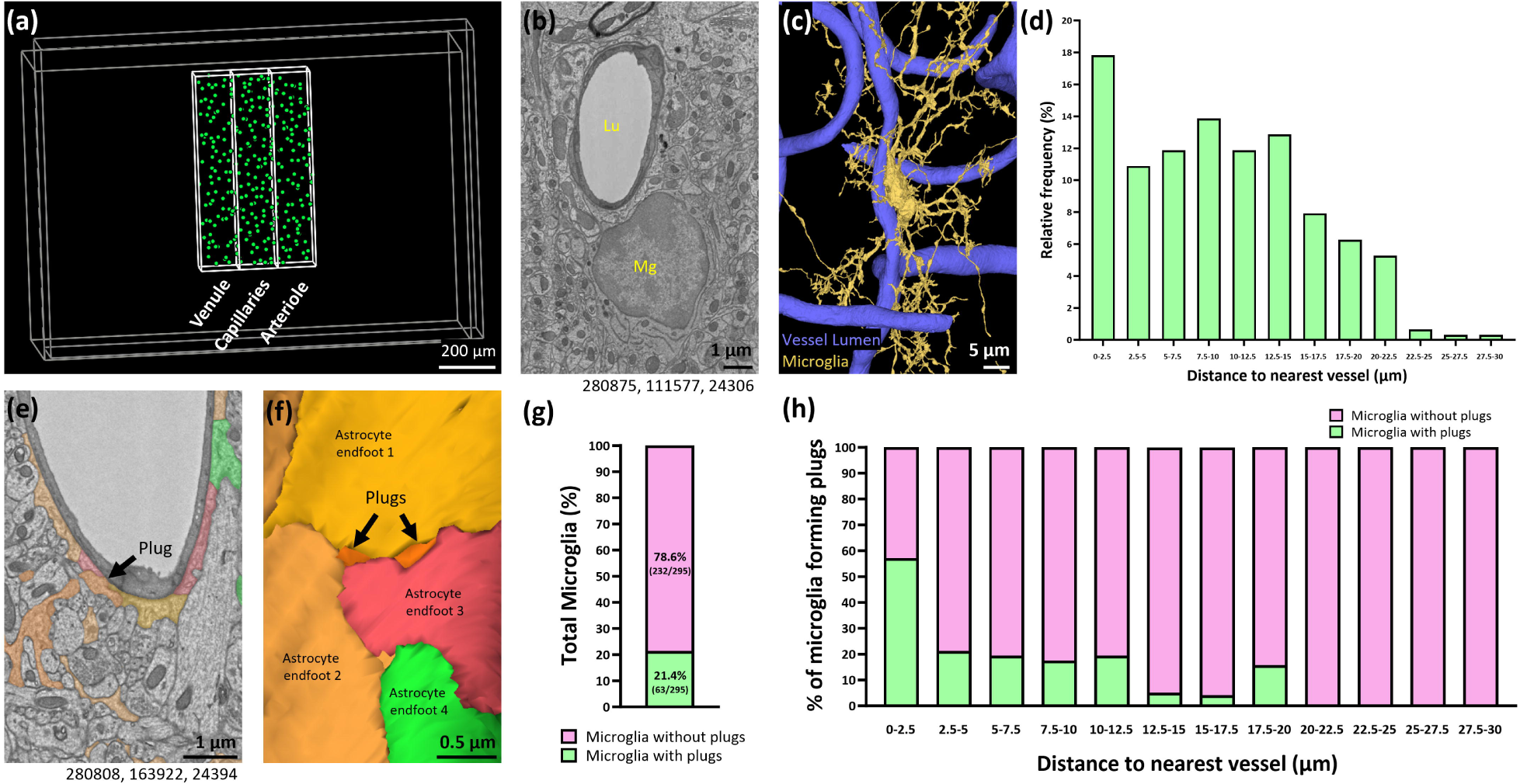
Microglia residing close to blood vessels are more likely to form plugs between astrocyte endfeet. (a) Annotations (green) showing the nuclear location of all microglia found in the venule, capillary and arteriole regions of interest. (b) Representative example of a microglia (Mg) in close proximity to a blood vessel (Lu = vessel lumen). (c) 3D reconstruction of (b). (d) The relative frequency of microglia binned by distance to vessel. (e) Representative example of a microglial plug between two separate astrocyte endfeet. (f) 3D reconstruction of (e). The plug in (e) is the plug on the right in (f). (g) The percentage of total microglia that form at least one plug. (h) The percentage of microglia that form at least one plug binned by the distance of their nuclear centre point to the nearest blood vessel. Numbers beneath figures (b) and (e) are xyz coordinates from the Cortical MM^3 dataset.

We found 21.4% of all microglia had at least one process directly contacting a blood vessel through gaps between adjacent astrocyte endfeet (Fig. 2(e-g)). We termed these contact points ‘plugs’. Of the microglia that form a plug, 57.1% were less than 2.5 µm from a blood vessel, but this percentage decreased the further microglia were away from the vasculature (Fig. 2(h)). This suggests the closer microglia are to the vasculature, the more likely they are to form plugs.

Microglia may also contact the vasculature directly by penetrating single astrocyte endfeet, which we define as ‘fenestrations’. We found very few convincing examples of these types of contacts. For example, we found one potential contact through a hole in an individual endfoot (Supplementary Fig. 1). However, for this and other examples, we were unable to conclusively say microglia were contacting the vessels through fenestrations, as the astrocyte endfoot may have simply become so thin that it is no longer discernible.^4^ Considering the lack of conclusive evidence that true fenestrations exist in astrocyte endfeet, we restricted our analysis to microglial plugs between endfeet, which provided more convincing evidence of the direct contact between microglia and the cerebral vasculature.

### Plugs are most frequently found close to microglia soma

Although microglia soma are generally smaller than somas of other cell types residing in the brain, their processes can reach >20 µm into the surrounding tissue. Given both microglia soma and their processes can spatially associate with the vasculature, this provides significant opportunities for plugs to form. We therefore next determined how many plugs microglia can form, and which parts of microglia most frequently form plugs.

The median number of plugs formed by plug-forming microglia was 2, and the most formed by a single microglia was 17 (Fig. 3(a-c)). Almost half of all plug-forming microglia formed just one plug (29/63, 46%), but 19% (12/63) formed five of more plugs (Fig. 3(d)). Of the 186 plugs found, 64% were formed <10 µm from the centre of a microglia nucleus (Fig. 3(e-g)). The furthest plug was 45.9 µm from its corresponding nucleus, but only 12.4% of plugs were >20 µm from the nucleus. It was also notable that all microglia with >5 plugs were within 5 µm of a blood vessel (Fig. 3(h)). Collectively, these results suggest plugs most commonly form near the microglia soma, and that the closer a microglia soma is to the vasculature, the more plugs it is likely to form.

**Figure 3:**
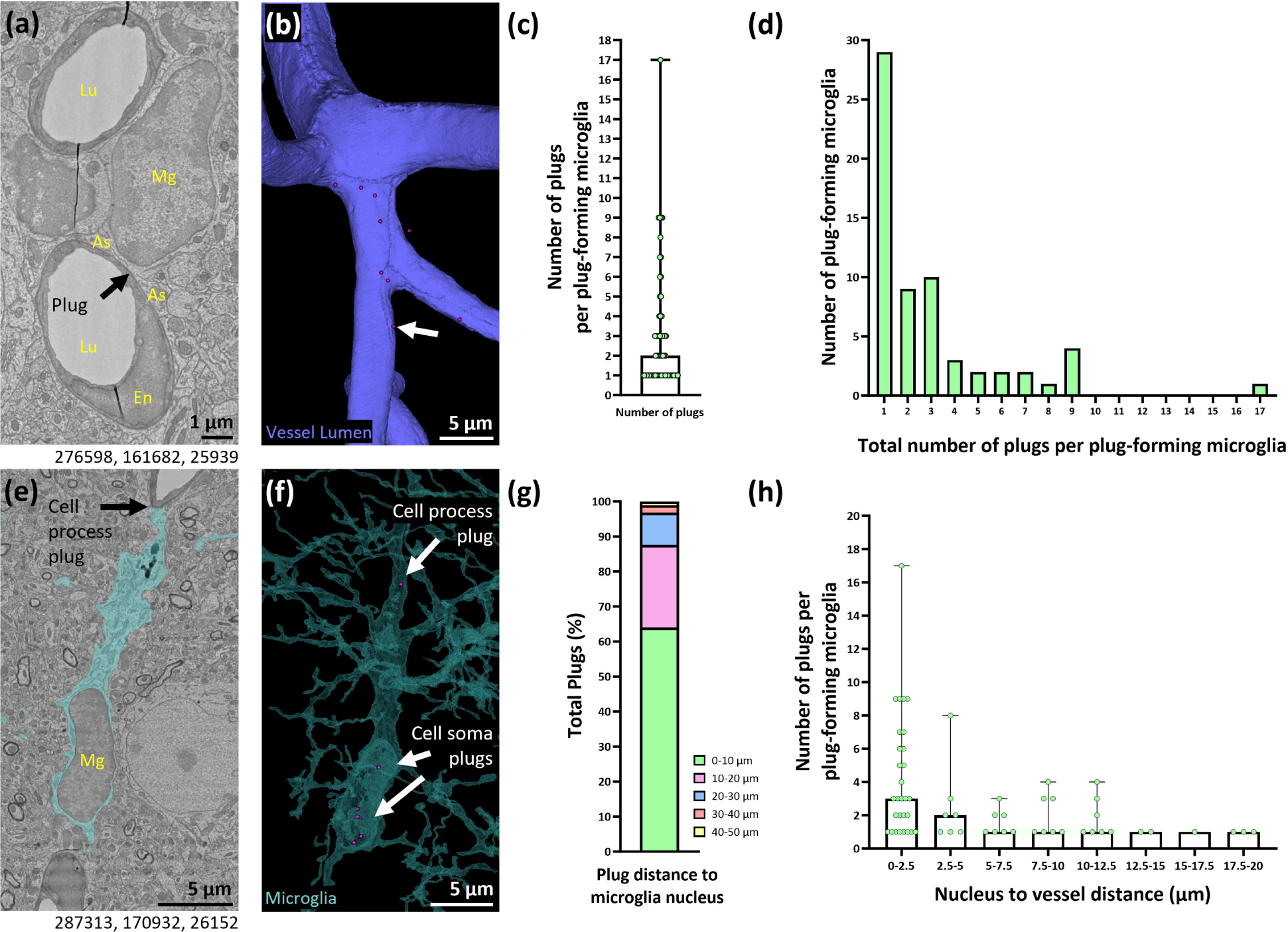
Plugs predominantly form near the microglial soma and microglia closer to the vasculature form more plugs. (a) Representative example of a plug-forming microglia (Mg) adjacent to a blood vessel (Lu = vessel lumen). As = astrocyte. (b) 3D reconstruction of the capillary lumen from (a) showing the location of plugs (magenta circles) formed by the microglia. (c) Quantification of the average number of plugs per plug-forming microglia (median + range). (d) Quantification of the number of plug-forming microglia that form between 1-17 plugs. (e) Representative example of a plug-forming microglia (Mg) with several plugs at the cell soma, and one plug formed at the process. (f) 3D reconstruction of microglia in (e) with the location of plugs (magenta circles) relative to the microglia soma and processes. (g) Quantification of the total percentage of plugs formed at various distances from the microglia nucleus. (h) Quantification of the number of plugs binned by the distance of plug-forming microglia to the nearest vessel (median ± range). Numbers beneath figures (a) and (e) are xyz coordinates from the Cortical MM^3 dataset.

### Microglial plugs are more prevalent at the venule end of the vascular tree

We next aimed to determine where on the vascular tree microglial plugs were most likely to form, which may provide clues about their function. We found no examples of plugs on either the arteriole or venule in our ROIs. Of the 186 plugs found, 70% formed on 1^st^ to 3^rd^ capillary branch orders away from the venule, with the number of plugs declining thereafter (Fig. 4(a-c) and Supplementary Fig. 2(a-b)). It is possible the higher prevalence of microglia <2.5 µm from a blood vessel in the venule ROI compared to the arteriole and capillary ROI (24.1% vs. 13.4% and 16.8% respectively, Supplementary Fig. 3(a-c)) may have influenced the total number of plugs near the venule. However, considering there were ∼8-fold more plugs in the venule ROI compared with the arteriolar ROI (Supplementary Fig. 2(b)), this seems unlikely to be the sole reason for the higher prevalence of plugs at the venule end of the vascular tree. Although we found plugs in the arteriole ROI, most plugs were closer in capillary branch order to the venule than the arteriole, therefore, for our dataset we show capillary order relative to the venule (Fig. 4(a-c)). Collectively, our data suggests plugs may be more common at the venule end of the vascular tree, rather than at the arteriolar end.

**Figure 4:**
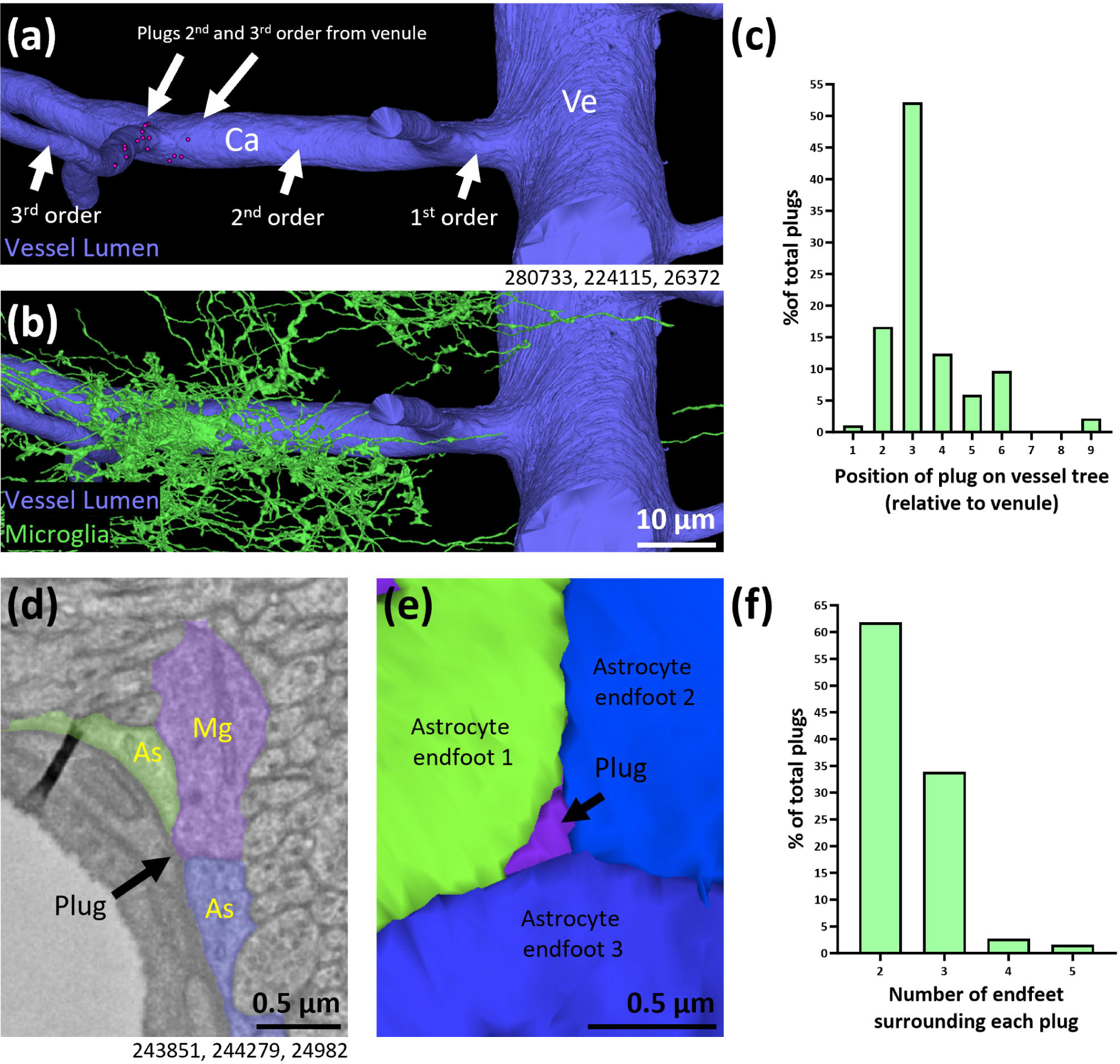
Microglial plugs predominantly form closer to the venule end of the vascular tree and are predominantly surrounded by 2 or 3 astrocyte endfeet. (a-b) 3D reconstruction of the microglia and vessel in Supplementary Fig. 2(a) with locations of plugs (magenta circles) on 2^nd^ and 3^rd^ order vessels away from a venule, without (a) and with (b) the microglia. (c) Quantification of the total percentage of microglial plugs at different levels of the vascular tree relative to the venule. (d) Representative example of a plug-forming microglia (Mg) between three astrocyte (As) endfeet. (e) 3D reconstruction of (d) showing the astrocyte endfeet with the microglial plug between them. (f) Quantification of the total percentage of microglial plugs with differing numbers of astrocyte endfeet surrounding them. Numbers beneath figures (a, d) are xyz coordinates from the Cortical MM^3 dataset.

### Microglial plugs are most commonly surrounded by two astrocyte endfeet

Astrocyte endfeet are predicted to form in a Voronoi-like tessellation pattern.^29^ Within such a pattern it is possible multiple endfeet can meet at the same point. We therefore next determined how many separate astrocyte endfeet were surrounding each of the plugs in our dataset. Microglial plugs were most commonly surrounded by two separate astrocyte endfeet (61.8%, Supplementary Fig. 4(a), Fig. 4(f)), but were also often surrounded by three endfeet (33.9%, Fig. 4(d-f)). We also found eight examples of plugs surrounded by as many as 4 or 5 endfeet, although these were rarer (2.7% and 1.6%, respectively, Supplementary Fig. 4(b)).

### Microglial plugs typically have a small surface area

Inspired by previous work which quantified endothelial and pericyte peg-and-socket size,^30^ we next determined the surface area of plug contact points on the vasculature to better understand plug diversity. The vast majority of plugs had a surface area <1 μm^2^ (95.2%), with only a small proportion >1 μm^2^ (4.8%, Fig. 5(a-g)). Further segregation of the data showed that of the plugs with a surface area <1 μm^2^, 83.6% of them were <0.1 μm^2^ in surface area (Fig. 5(h)). The small number of plugs that were ≥1 μm^2^ ranged in size from 1.3 μm^2^ - 14.3 μm^2^ (Fig. 5(d-f, i)), illustrating they can vary hugely in size, but that the majority are relatively small (<0.1 μm^2^) in surface area.

**Figure 5:**
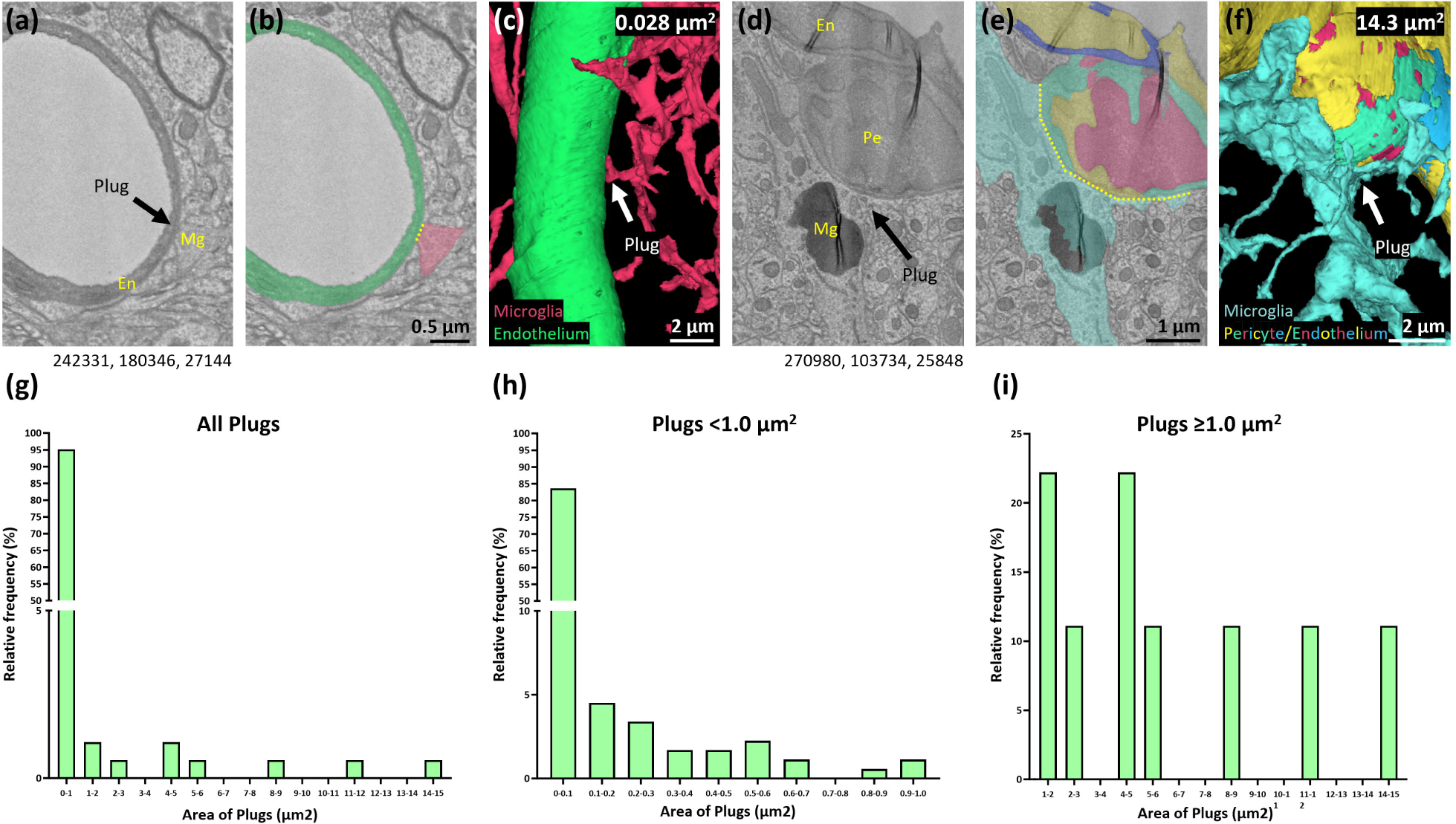
Microglial plugs exhibit a large range of surface areas, but the majority are <0.1 μm. (a) Representative example of a microglial (Mg) plug with a small surface area, on an endothelial cell (En). (b) Segmentation of the microglia and the endothelium from (a). (c) 3D reconstruction of the microglia and the endothelium from (a). (d) Representative example of a microglial (Mg) plug with a large surface area, on a pericyte (Pe). (e) Segmentation of the microglia and pericyte from (d). (f) 3D reconstruction of the microglia and pericyte from (d). Yellow dotted lines indicate the width of the contact point in a single z-plane. (g) Quantification of the relative frequency of the surface area of all microglial plugs. (h) Quantification of the relative frequency of microglial plugs <1.0 μm in surface area. (i) Quantification of the relative frequency of microglial plugs ≥1.0 μm in surface area. Numbers beneath the figures (a) and (d) are xyz coordinates from the Cortical MM^3 dataset.

### Microglia plugs may be transient

It is understood that microglia processes are constantly surveying their microenvironment.^31, 32^ Microglia soma can also shift over the course of several hours or weeks,^32–34^ with many of these movements associated with blood vessels.^34^ To illustrate that microglia-vascular contacts may be transient, we used *in vivo* two-photon microscopy to image microglia soma and processes spatially associated with pericytes on blood vessels at the arteriole and venule ends of the vascular tree, and within the capillary bed, through cranial windows implanted in NG2DsRed×CX3CR1^+/GFP^ mice. Microglia (CX3CR1^+/GFP^) and pericytes (NG2DsRed) were repetitively imaged over 7-minute periods. These images revealed microglia processes actively extending and retracting toward pericyte soma and processes on the cerebral vasculature over a brief timeframe (Fig. 6(a), Video 1). One caveat to this is that many of the plugs we identified in the EM dataset are <1 μm^2^ and may therefore be beyond the limit of detection using two-photon microscopy. However, these results highlight how microglia-vessel plugs may be transient, but we cannot rule out that some contacts may be long-lasting.

**Figure 6:**
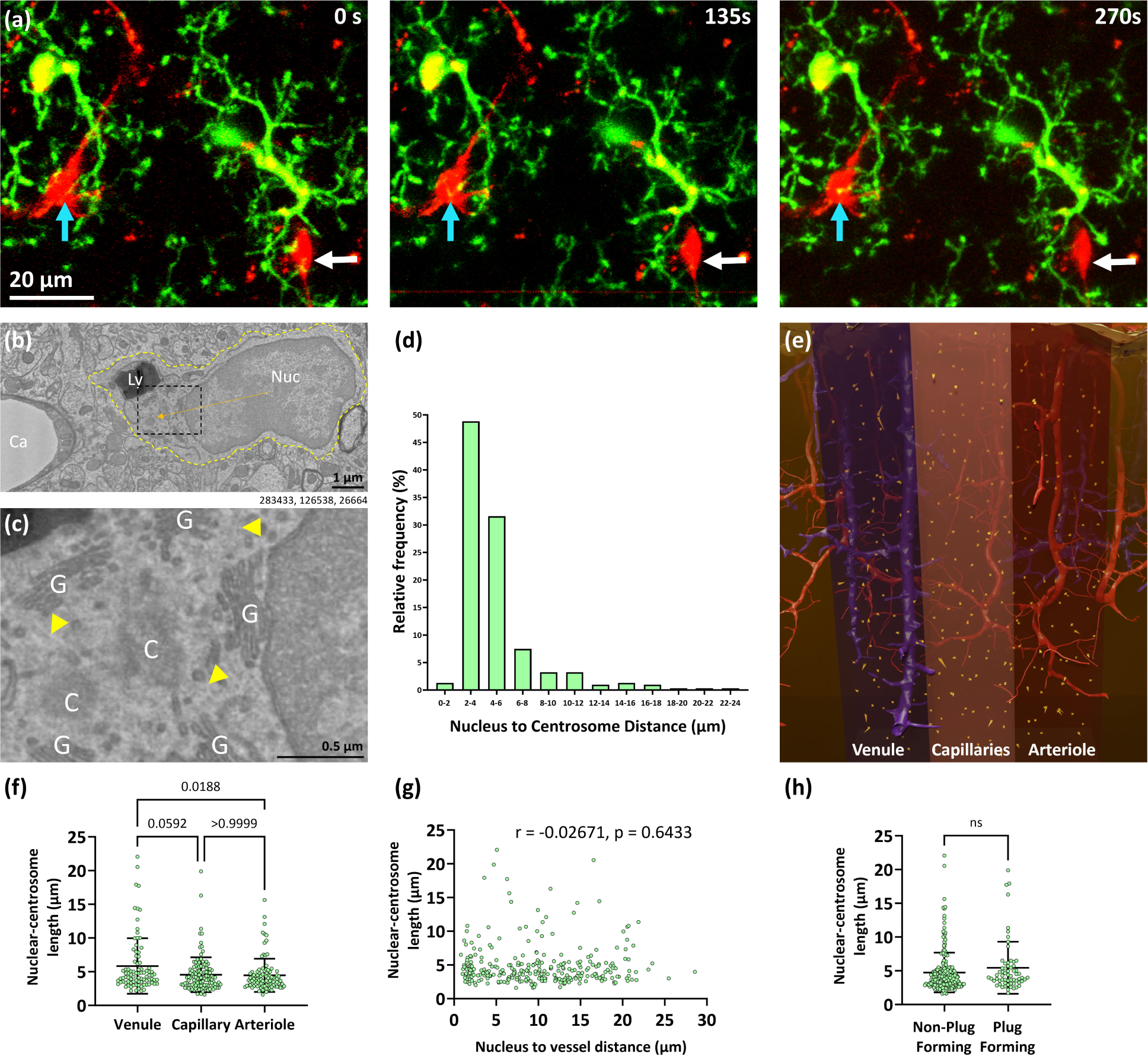
Microglial plugs may be transient. (a) 231μm thick projection images of NG2DsRed-positive pericytes (red) and CX CR1^+/GFP^-positive microglia (green) in layers II/III of the somatosensory cortex of an adult female NG2DsRed×CX CR1^+/GFP^ mouse imaged using *in vivo* two-photon microscopy. Cyan arrows indicate a microglia process moving toward a pericyte soma over time. White arrows indicate a microglia process moving away from a pericyte soma over time. (b) Representative example of a microglia (Nuc = nucleus, Ly = lysosome, yellow dashed outline = outline of the cell) near a capillary (Ca). The orange arrow indicates the NC length. (c) Enlargement of dashed black square in panel (b). Microglial centrioles (C) are surrounded by the golgi network (G). Microtubules (yellow arrowheads marking faint lines) radiate from the centrioles. (d) Quantification of the relative frequency of different nuclear-centrosome lengths for all microglial in the dataset. (e) Schematic view of the Cortical MM^3 volume highlighting the regions of interest we analysed and the nuclear-centrosome lengths of microglia (orange arrows) within these regions. The shaded areas denote the different regions analysed. See Supplementary Fig. 5 for wider top-down and side views of where the regions of interest are in the volume. (f) Quantification of NC length of microglia in the venule, capillary and arteriole regions (analysed with a Kruskal-Wallis test; Kruskal-Wallis statistic = 8.442, p = 0.0147), with Dunn’s post hoc test (mean ± SD). (g) Correlation of NC length and microglial nucleus to the edge of the nearest vessel length (Spearman correlation). (h) Quantification of NC length in non-plug-forming and plug-forming microglia (mean ± SD). Numbers beneath figure (b) are xyz coordinates from the Cortical MM^3 dataset.

Unfortunately, microglia soma and process motility cannot be inferred from static EM images. However, the orientation of the nuclear-centrosomal (NC) axis may provide a proxy metric for the direction cells may be migrating in, as the centrosome is thought to be located between the leading edge of migration and the nucleus in several cell types, with the nucleus shifting toward the back of the migrating cell.^35–37^ This axis has been used to assess microglia migration *in vitro*^35^ and the movement of the centrosome into one branch of a microglia can predict microglia phagocytosis in zebrafish *in vivo.* ^38^ We found the location of the centrosome in each microglia (Fig. 6(b-c)) to determine the NC distance. The average NC distance was 4.9 ± 3.1 μm (Fig. 6(d)). In microglia with longer NC lengths (i.e. >7 μm), the centrosome was generally positioned in a large process emanating from the cell body (see Video 2), which may suggest these processes are moving toward a target (see Supplementary Table 3 for all centriole coordinates). In our dataset, microglia within the venule region had longer NC lengths than those in capillary and arteriole regions (5.8 ± 4.1, 4.6 ± 2.6 and 4.5 ± 2.5 μm, respectively, Fig. 6(d)). However, there was no correlation between NC length and proximity of microglia nuclei to vessels (Fig. 6(f)). Furthermore, we found no difference in the NC length between plug- and non-plug-forming microglia (5.5 ± 3.9 vs. 4.7 ± 2.9 μm, Fig. 6(h)). If NC length is considered a proxy of microglial motility, or movement toward a target, our data suggests there is no clear evidence plug-forming microglia are moving differently to non-plug forming microglia.

### Microglia plugs contact both endothelial cells and pericytes

Due to their close spatial association with blood vessels, microglia may contact different cellular elements of the vasculature. Their location on the vascular tree can dictate the types of cells microglia may come into contact with. For example, pericytes are present throughout the capillary tree in the brain. Furthermore, perivascular fibroblasts (PVFs) and perivascular macrophages (PVMs) are found on large ascending venules, penetrating arterioles, and first-order capillaries, with some processes reaching third-order capillaries, but are rarely found elsewhere on the vascular tree.^4, 27^

In our dataset, the most frequently contacted cells by microglial plugs were endothelial cells (69%, Fig. 7(a-c)), followed by pericytes (24%, Fig. 7(c-e)), with some microglial plugs contacting both endothelial cells and pericytes at the same time (7%). We did not find any plugs contacting either PVFs or PVMs (see Supplementary Table 2 for location of PVFs and PVMs in our ROI). We further probed the dataset to determine where plugs on endothelial cells and pericytes were most prevalent. Most plugs were directly on endothelial cell (62%, Fig. 7(b, f)), and pericyte (23%, Fig. 7(e, f)) processes. There were occasional contacts with both endothelial soma (5%, Fig. 7(a, f)) and pericyte soma (1%, Fig. 7(d, f)), and some plugs contacted multiple cellular elements at the same time (Fig. 7(f)).

**Figure 7:**
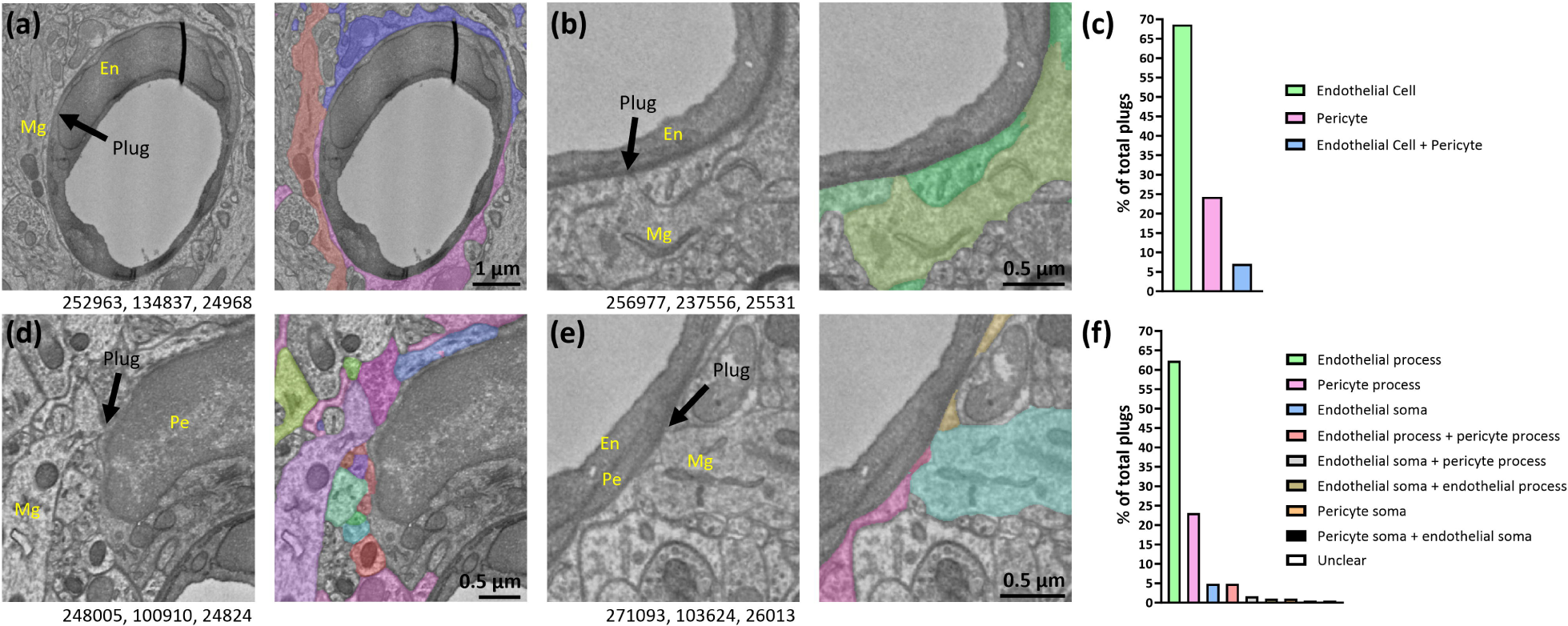
Microglial plugs contact both endothelial cells and pericytes. (a) Representative example of a microglia (Mg) contacting an endothelial cell nucleus (En). Image to the right shows the astrocyte endfeet and microglial plug, in colour. (b) Representative example of a microglia contacting an endothelial process. Image to the right shows the astrocyte endfeet and microglial plug, in colour. (c) Quantification of the percentage of all microglial plugs that contact endothelial cells and pericytes. (d) Representative example of a microglia contacting a pericyte (Pe) soma. Image to the right shows the astrocyte endfeet and microglial plug, and several plugs from other cell types, in colour. (e) Representative example of a microglia contacting a pericyte process. Image to the right shows the astrocyte endfeet and microglial plug, in colour. (f) Quantification of the percentage of all microglial plugs contacting different locations on each cell (e.g. soma or processes) of endothelial cells or pericytes. Numbers beneath the figures are xyz coordinates from the Cortical MM^3 dataset for each representative image.

## Discussion

This study used the largest publicly available 3D EM dataset of a mouse brain to investigate the ultrastructural relationship between microglia and the cerebral vasculature. Our results show microglia have direct contact with the brain vasculature through gaps between adjacent astrocyte endfeet, termed “plugs”. Our findings have implications for the physiological function of microglia at the vasculature, and for the crosstalk between parenchymal brain cells and cells of the vasculature in health and disease.

Consistent with previous reports in the brain^12, 13, 15, 19^ and retina,^40^ we observed a close spatial relationship between microglia and the vasculature in the Cortical MM^3 dataset. Over 50% of microglia were found residing within 10 μm of the vasculature, and ∼18% of microglia were within 2.5 μm of the vasculature. The furthest distance between the centre of a microglia soma and a vessel edge was 28.6 μm, which fits with reported average inter-capillary distances of ∼40-60 μm in mammalian cortices.^41, 42^ Considering microglia were 9.7 μm from capillaries on average, this suggests the majority of microglia prefer to reside spatially closer to vessels than away from them. Despite this close spatial relationship, only ∼1 in 5 microglia formed plugs with the vasculature through gaps between adjacent astrocyte endfeet, suggesting most microglia are not actively contacting vessels at any given time.

Although we restricted our analysis to microglia-vessel contacts that occurred between adjoining astrocyte endfeet, we cannot rule out that microglia may contact blood vessels through holes in individual astrocyte endfeet (contacts we term ‘fenestrations’). We probed the dataset for examples of such contacts but found little convincing evidence that they exist. The primary reason for doubt is that astrocyte endfeet can become so thin that they blend into the surrounding plasma and basement membranes, making it impossible to determine if there is truly a lack of an endfoot.

In our study we found no microglial plugs on the venule or arteriole, only on capillaries. Evidence suggests arterioles have the thickest endfeet dimensions surrounding them, followed by venules and then capillaries.^3, 43^ It is possible plugs are present on capillaries rather than arterioles and venules due to thicker astrocyte endfeet inhibiting plug formation. One of the striking findings in this work was that plugs were more prevalent on capillaries toward the venule end of the vascular tree, raising intriguing functional implications due to the differing functions of blood vessels across the vascular tree. Nanoparticles have been shown to enter the brain via post-capillary venules that project onto ascending venules,^44^ which is consistent with the location where immune cells may traffic into the brain.^45^ The prevalence of plugs at the venule end of the tree, on capillary orders often referred to as post-capillary venules,^44^ suggest microglia may functionally interact with immune cells at these locations, a hypothesis worth exploring in future studies.

We do not know the significance of our finding that plugs commonly occur at inter-endfoot junctions containing 2-3 endfeet. It is possible this may be governed by the relative frequency of such junctions (i.e. 2 or 3 endfeet at a junction might be the most common occurrence). In support of this theory, we have previously found that astrocyte endfeet form sharp lines between them in a Voronoi-like pattern, which means most junctions will be between 2 or 3 astrocyte endfeet.^4^

It was notable that 95% of microglial plugs covered <1 μm^2^ in surface area. However, 3.8% of plugs were >2 μm^2^ and appeared to essentially take the place of astrocyte endfeet over large stretches of the vasculature. For biological context, proteins as large as 500 kDa have a predicted radius of ∼5 nm.^46^ Extracellular vesicles capable of carrying proteins, lipids and nucleic acids can range in diameter from as small as 30 nm to as large as 4 μm.^47^ The smallest plug in our dataset had a diameter at the centre z-plane of ∼40 nm and the median diameter of all our plugs was ∼160 nm. Collectively, this data suggests microglial plugs have biological relevance as they are large enough to enable the passage of small to large extracellular vesicles between microglia and cells of the vasculature.

We do not yet know how dynamic microglial plugs are. It is possible they are constantly changing size and shape and that plug formation is dynamic, a theory supported by the many examples we found of microglial processes nearby inter-endfeet junctions. It is also possible inter-endfeet gaps grow and shrink as recently suggested by others,^3^ adding another layer of complexity to the permanence or transience of plugs. Our *in vivo* two photon videos highlight how dynamic microglia processes are around capillaries even over short 7-minutes periods. However, it is equally plausible some contacts may be more permanent, or that plug formation occurs regularly at specific ‘hotspots’. Studying plug dynamics using *in vivo* two photon microscopy will be an important future direction, but it is imperative to highlight that many of the smaller plugs are likely below the limits of resolution of two photon microscopes, which is ∼500 nm in xy and ∼2-3 um in z. Therefore, ultrastructural analyses will remain important to better define how microglial plugs change in different physiological and pathological conditions.

A proportion of microglia soma are actively migrating at any given time, with one previous study suggesting ∼5% of microglia soma shift 1-2 μm over the course of 10h.^32^ Other studies suggest ∼14-16% of microglia soma shift >5 μm over the course of days to several weeks.^33, 34^ Interestingly, as many as half of these migratory movements may be associated with blood vessels.^34^ Determining migration in static EM images is clearly challenging, however evidence suggests the nuclear-centrosome (NC) axis may provide a surrogate of cell migration in many cells, including microglia.^35, 38, 39^ Although a role for the centrosome in microglia migration has not been definitively established, there is one report that microglia centrosomes can move up to 2 µm/min toward apoptotic neurons in the zebrafish optic tectum 3 days post-fertilisation *in vivo*.^38^ In contrast, there is some evidence centrioles are in the central part of the cell when macrophages migrate *in vitro* and therefore might not be connected to migration^39^, however this evidence is from 2D cultured cells that migrate without significant tissue resistance. Because plug formation by microglia could be an active process, we were interested in assessing if the NC length differed in plug and non-plug forming microglia. However, we found no evidence to support a difference in NC length between plug and non-plug forming microglia. This suggests that if NC length is a proxy marker of microglia motility, then there is no significant difference in motility between plug and non-plug forming microglia.

As the vasculature is surrounded by the basement membrane,^48^ any contact plugs make with cells of the vasculature also involve contact with the basement membrane. The thickness of the basement membrane is difficult to determine at sites of microglia-vascular contact as it blends into the plasma membrane of both microglia and cells of the vasculature in the EM dataset. Pericytes, astrocytes and endothelial cells bind to the basement membrane through integrin or dystroglycan receptors,^48^ but it is not clear if microglia are also capable of doing so. There is however some evidence microglia can form tight junctions with endothelial cells during inflammation, but it is not clear if this also occurs in normal physiology.^16^

Following identification of plugs, we found that the most common contacts were with endothelial cells. However, we also saw a significant number of contacts at pericytes, both at pericyte processes and pericyte soma, consistent with previous reports of spatial associations between microglia and pericytes.^12, 15, 40, 49^ Microglia plugs were ∼3-fold more common at endothelial cells than pericytes, which may be explained by the relative ratio of endothelial cells to pericytes in the brain. Although it is not fully clear exactly how much of the cerebral vascular surface is covered by pericytes, some estimates suggest ∼22-30%^50^, although others suggest higher coverage^51^. It is also worth noting pericyte coverage is likely higher at the arteriole end of the capillary tree where ensheathing pericytes are present,^52^ compared to the venule end of the capillary tree which contains thin-strand and mesh pericytes. This suggests that if the frequency of plug contact with endothelial cells and pericytes is the same as their relative ratio along the vascular tree, then microglial plugs have no clear preference for endothelial cells or pericytes.

The contacts we observed with pericyte soma were particularly intriguing. There were three plugs contacting pericyte soma and it was notable that at all three locations, the pericytes were also contacted by plugs from other cell types (see Fig. 7(d) for an example). These observations suggest some pericyte soma may have a specific lack of astrocyte endfoot coverage, allowing parenchymal cells access to pericytes without the structural interference of astrocytes. This finding is in agreement with a previous report suggesting the endfoot covering of pericytes may be incomplete, in contrast to the covering of endothelial cells.^2^

Microglia plugs may have many functions. It has recently been suggested microglia may play roles in the regulation of blood flow.^13, 15^ Furthermore, microglia have been identified as a major source of platelet derived growth factor B (PDGF-B),^53^ a key molecule for pericyte homeostasis^54^ and BBB integrity.^54^ Considering these findings, it is attractive to suggest plugs may serve as an area through which microglia can release chemical mediators to influence pericytes and endothelial cells, thereby indirectly influencing vascular function. However, it is also possible some chemical mediators can be released from microglia and diffuse across astrocyte endfeet,^55, 56^ which would bypass the need for direct microglia-vessel contact.

Plugs may also allow microglia to closely communicate with immune cells which may traffic into the brain at the same areas these plugs appear to be most prevalent (the venule end of the vascular tree). Furthermore, they may play a role in water homeostasis and waste removal in and out of the brain via the glymphatic system,^57^ by plugging gaps where fluid flow could occur. Alternatively, microglia may release antigens into the glymphatic system through these plugs. One caveat to these ideas however is that it is not yet known if glymphatic flow occurs around capillaries, having predominantly been studied in larger vessels.

Our study has several limitations. The primary limitation is that our dataset is derived from one region of a single mouse brain that has undergone a cranial window implantation and two-photon imaging prior to tissue collection and electron microscopy, so therefore it may not fully reflect normal physiology.^58^ Furthermore, a recent study has suggested different methods of tissue processing in preparation for EM imaging can affect astrocyte endfoot coverage of vessels, which if true would likely change the number or size of microglial plugs observed.^59^ However, the reviewers of that paper raised some important concerns about their cryo-fixation method. Furthermore, it is likely some plugs were missed as the segmentation, which aids the analysis, is not always accurate. As a result, our quantification may have underestimated the prevalence of microglia-vascular contacts. It is also important to note plugs may not be restricted to microglia, as we observed several examples of other cells also forming plugs between astrocyte endfeet, however we have not formally analysed the cellular origin of these other plugs.

In summary, although astrocyte endfeet cover almost the entire cerebral vasculature,^2^ this study has established the existence of microglia plugs directly contacting blood vessels between adjacent astrocyte endfeet. Our work provides a platform to begin understanding the formation and dynamics of microglial plugs, as well as their function in brain health and disease.

## Acknowledgements

We thank Jo-Maree Courtney for assistance with excel formula and functions used for data analysis. We thank Bill Bennett and Alison Canty for technical assistance and advice with two-photon microscopy imaging. We thank Jenna Ziebell for implanting the cranial windows used to create the videos in this study. We thank Andy Shih and Stephanie Bonney for kindly providing the location of the PVFs in our dataset. We acknowledge and thank the MICrONS Consortium for producing the electron microscopy data used in this study and for making it freely available to the international research community.

## Author Contribution Statement

GPM: conceptualization, methodology, software, validation, formal analysis, investigation, data curation, writing – original draft, review and editing, visualization, supervision, project administration. CGF: conceptualization, methodology, software, investigation, writing – review and editing. BAS: conceptualisation, methodology, resources, writing – original draft, review and editing, supervision, project administration, funding acquisition. SG: conceptualization, methodology, software, validation, formal analysis, investigation, resources, data curation, writing – original draft, review and editing, visualization, supervision

## Supplementary material

Supplementary material for this article is available online. All data can be provided by the corresponding authors upon reasonable request.

## Funding statement

This research was supported by an NHMRC Boosting Dementia Fellowships (APP1137776, BAS) and an NHMRC project grant (APP1163384, BAS). The funding sources had no role in the study design, in the collection, analysis and interpretation of the data, in writing the report, or in the decision to submit the article for publication.

## Conflict of interest disclosure

No conflicts of interest to disclose.

## Notes

### Competing Interest Statement

The authors have declared no competing interest.

