## Supplementary Materials for "Microglia contact cerebral vasculature through gaps between astrocyte endfeet"

**Author affiliations**

Dr. Søren Grubb

Full address: Blegdamsvej 3B, Copenhagen N, Denmark, DK-2200, Ph: +45 27290007,

**Or:** Dr. Gary Morris

Full address: Medical Sciences Precinct, 17 Liverpool Street, Hobart, TAS 7000, Australia, Ph: +61 3 6226 4851;

### Video legends

Video 1 – Showing in vivo two-photon microscopy data of NG2DsRed × CX3CR1^+/GFP^ mice near a penetrating arteriole, near the capillaries and near an ascending venule. Orange arrows indicate locations of pericyte somas. Length of each recording is 7 min and 30 s. The volume and surface renderings were done using Imaris software.

Video 2 – Showing vascular cells and a microglia with a main process partly covering a pericyte soma protruding through a gap between astrocyte endfeet. The microglia has two large lysosomes in the main process where the centrosome is also located. The main process is also invaginated by a primary cilium of a nearby interneuron. The volume rendering and segmentation was done using Amira software.

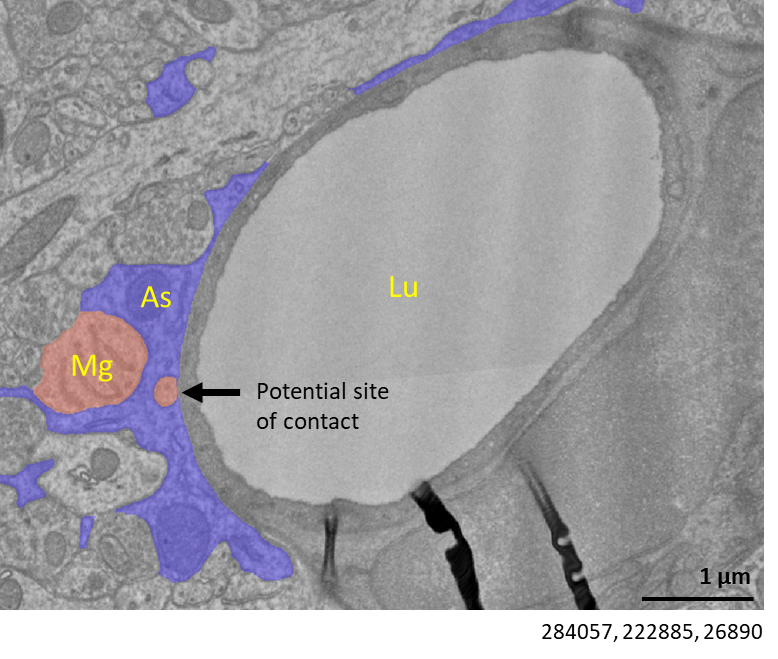

**Supplementary Figure 1:**

Example of potential capillary contact (Lu = vessel lumen) by a microglia (Mg) through a fenestration in a single astrocyte (As) endfoot. Numbers beneath figure are xyz coordinates from the Cortical MM^3 dataset.

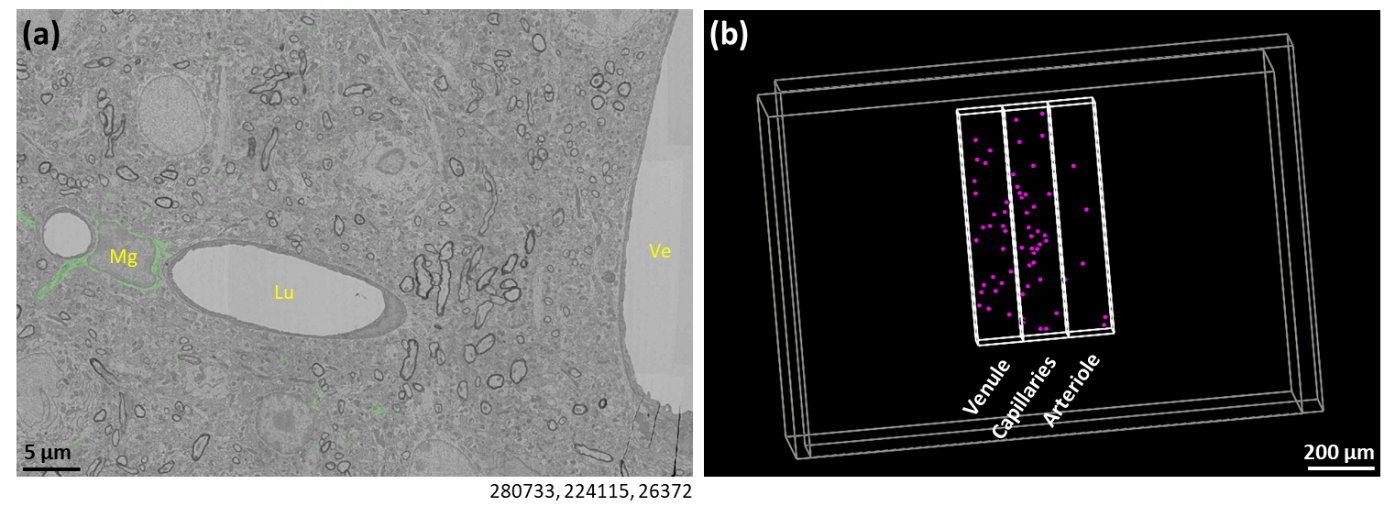

**Supplementary Figure 2:**

(a) Representative example of a plug-forming microglia (Mg) forming plugs on 2^nd^ and 3^rd^ order vessels (Lu = vessel lumen) away from a venule (Ve) (see Fig. 4(a-b) for plugs formed by this microglia). Numbers beneath figure are xyz coordinates from the Cortical MM^3 dataset. (b) Location of all plugs (magenta circles) in the venule, capillary and arteriole regions of interest.

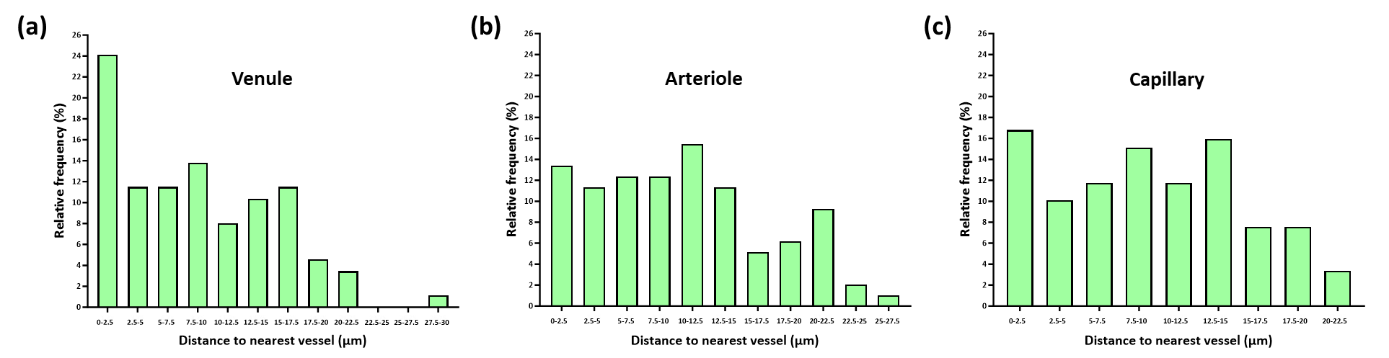

**Supplementary Figure 3: Microglia proximity to cerebral blood vessels in the venule, arteriole and capillary regions**

(a-c) The relative frequency of microglia binned by distance to vessel in the (a) venule (n=87), (b) arteriole (n=97) and (c) capillary (n=119) regions.

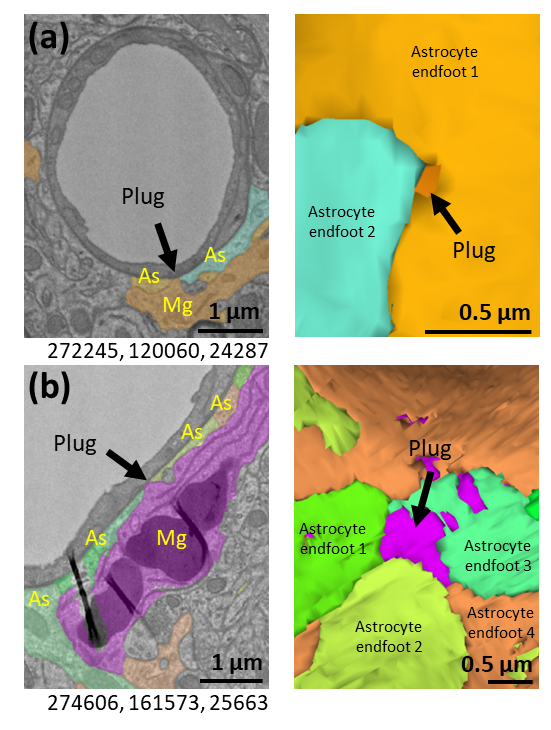

**Supplementary Figure 4:**

(a-b) Representative example of plug-forming microglia (Mg) between (a) two and (b) four astrocyte (As) endfeet. The right images show 3D reconstructions of (a) and (b) with astrocyte endfeet surrounding the microglial plugs. Numbers beneath figures (a-b) are xyz coordinates from the Cortical MM^3 dataset.

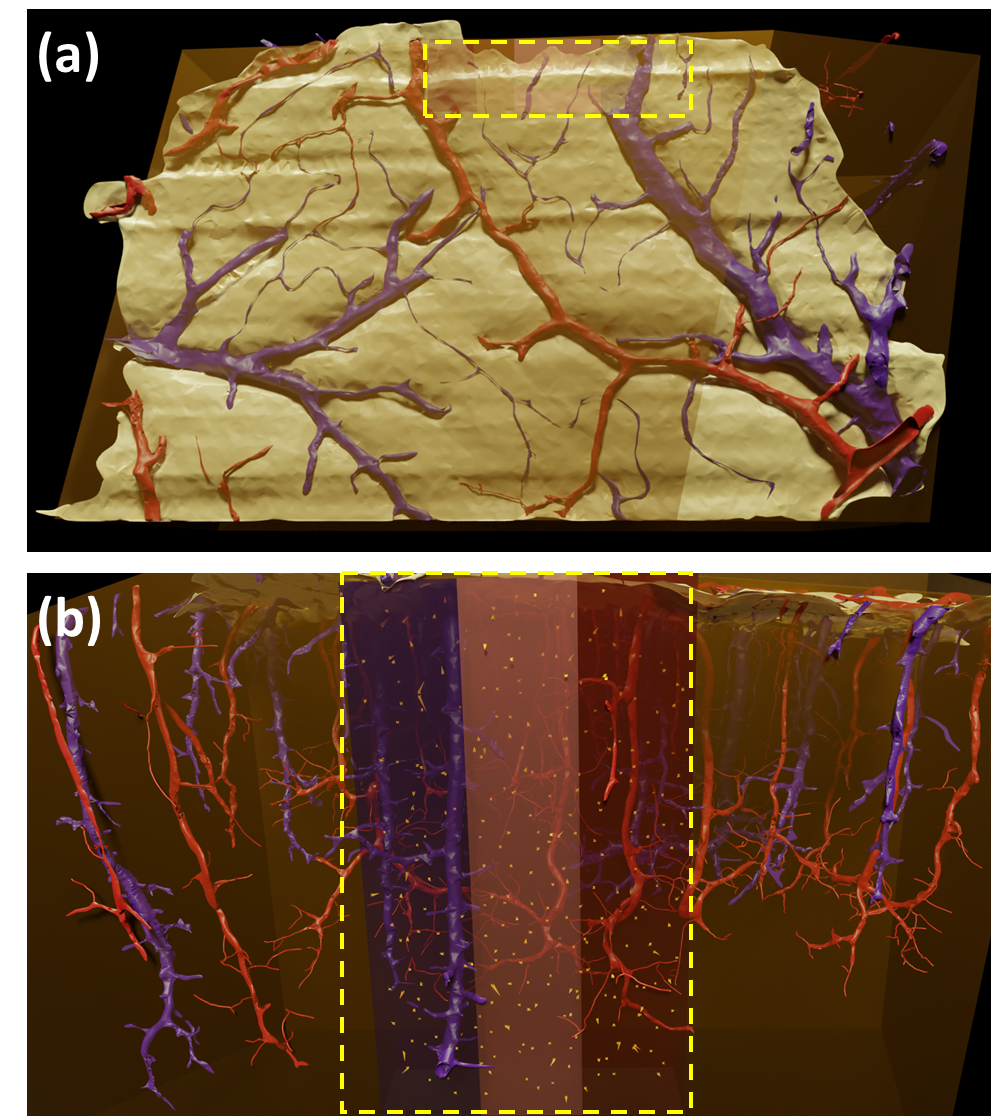

**Supplementary Figure 5:**

(a) Schematic of the top of the Cortical MM^3 volume dataset showing pial arterioles in red and pial venules in purple. Yellow dashed box indicates the position of our regions of interest. Transparent orange box indicates limits of the dataset. (b) Schematic view of the Cortical MM^3 volume highlighting the regions of interest we analysed (yellow dashed box) and the nuclear-centrosome lengths of microglia (orange arrows) with these regions. See Fig. 6(d) for a cropped version of this figure.
